# Sex and dominance status affect allogrooming in free-ranging feral cattle

**DOI:** 10.1101/2023.05.08.539791

**Authors:** George M. W. Hodgson, Kate J. Flay, Tania A. Perroux, Wai Yan Chan, Alan G. McElligott

## Abstract

Social interactions are fundamental properties of gregarious species, helping to establish dominance hierarchies and maintain social bonds within groups, thus having significant effects on fitness. Cattle (*Bos taurus*) are social ungulates which engage in affiliative and agonistic relationships with other individuals. Although there are approximately 1.5 billion cattle on the planet, the opportunity to research cattle behaviour in free-ranging groups is rare, as there are few feral populations worldwide. Cattle engage in positive social behaviours such as allogrooming, where one individual licks the body of another. The relationship between affiliative behaviours and other individual characteristics (such as sex and dominance status) are frequently studied in other gregarious species, but are largely undetermined in cattle. To investigate the relationships between sex, dominance status and allogrooming, we observed a mixed-sex feral cattle herd in Hong Kong, recording dominance interactions and allogrooming events. We found that dominant females received more allogrooming than subordinate females, but subordinate females did not perform more than dominant ones. Males performed allogrooming more towards females than other males, but females groomed both sexes equally. Sex affected dominance position, with males obtaining higher status than females, but not all females were subordinate to males. These preferential allogrooming patterns improve our knowledge of sex-specific interactions, and help us to understand the dynamics of agonistic and affiliative behaviours in multi-male, multi-female ungulate groups. Studying a free-ranging feral population provides us with a unique insight into ungulate behavioural patterns and the evolution of cattle social behaviours.

## INTRODUCTION

Social behaviours help to establish bonds between partners, maintain stable hierarchies within groups, and are critical for the fitness of individuals (Hobson et al., 2021; Sussman et al., 2005). Interactions among group-living individuals are key aspects of animal social structures and allow us to examine patterns of sociality (Rubenstein & Abbot, 2017; Silk, 2007). Minimising predation risk is a critical benefit of group-living, conferring advantages to all group members (Ward & Webster, 2016). The degree of social organisation affects lifespan and survival, with group-living mammalian species living longer than solitary species (Zhu et al., 2023) and strong affiliative bonds reducing the risk of injury (Pavez-Fox et al., 2022). The social organisation of a species also influences the distribution of parasites within a community, and affects the vulnerability of a mammalian population to infectious disease (Altizer et al., 2003). The strength and pattern of these social bonds can be measured by repeated affiliative social interactions between individuals, such as allogrooming (Silk et al., 2013).

Allogrooming, also known as social grooming, involves the acting individual (performer) cleaning the body of another individual (receiver), often directing this behaviour towards skin and fur blemishes (Dunbar, 1991; Schino, 2007). Allogrooming has hygienic, social and physiological effects (Laister et al., 2011; Pfoh et al., 2021; van Dierendonck et al., 2009). The basic foundations for allogrooming behaviour have been established through extensive non-human primate research, focusing on understanding the evolution of sociality by examining dyadic interactions (Di Bitetti, 1997; Lehmann et al., 2007; Schino et al., 1988). One of the primary functions of allogrooming is ectoparasite removal (Akinyi et al., 2013; Dunbar, 1991), with allogrooming directed towards inaccessible body regions that the receiver cannot reach via self-grooming (King & Gurnell, 2019; Pfoh et al., 2021). Additionally, performing and receiving grooming provides physiological benefits for both partners (Schino et al., 1988), such as tension reduction (Laister et al., 2011; Yates et al., 2022). Allogrooming also serves a social purpose by promoting and maintaining social bonds between individuals (Schino, 2001; Schino & Aureli, 2009; Silk et al., 2003).

The distribution of allogrooming within a group is not random, and can be affected by individual characteristics, such as dominance status and sex (de Waal, 1997). For example, allogrooming can be directed towards higher-ranking individuals for exchanging rank-related services (Gareta García et al., 2021; Gumert, 2007; Schweinfurth & Taborsky, 2018). However, allogrooming may also be directed towards unfamiliar or subordinate individuals when used to strengthen social bonds and increase group cohesion (Di Bitetti, 1997; Fedurek & Dunbar, 2009; Šárová et al., 2016). Sexually-dimorphic allogrooming can occur when males gain specific benefits from females, such as exchanging allogrooming for mating opportunities (Barelli et al., 2011; Gumert, 2007).

Repeated agonistic interactions between individuals play a major role in the formation of dominance relationships, and can lead to the establishment of a stable hierarchy (Dehnen et al., 2022; Drews, 1993; Krahn et al., 2022). This allows individuals to evaluate their chances of winning a conflict, avoid injury risk and avoid unnecessary costs of time or energy loss by limiting conflict escalation (Holekamp & Strauss, 2016). An individual’s position in the hierarchy affects its resource access, reproductive success, and health, with higher dominance status expected to confer net fitness benefits (Archie et al., 2012; Bateman-Neubert et al., 2023; Blomquist et al., 2011; Majolo et al., 2012). Assessing the hierarchy of a population allows further examination of sociality, as dominance also affects movement leadership and the distribution of affiliation and aggression within the group (Hobson et al., 2021; Šárová et al., 2010).

Cattle (*Bos taurus*) are highly social animals and engage in affiliative and agonistic behaviours (Krahn et al., 2022; Marino & Allen, 2017; Padilla de la Torre & McElligott, 2017). Variation in social and non-social behaviours has been noted between European taurine breeds (*Bos taurus taurus*) and the indicine breeds (*Bos taurus indicus*; Hubbard et al., 2021), and differences between populations may be dependent on cattle breed and sub-species (Sartori et al., 2016; Solano et al., 2005). Individual characteristics (such as sex, age, and dominance status) are expected to affect the rates of social behaviours and allogrooming (Hubbard et al., 2021; Šárová et al., 2013), but the relationship between individual dominance status and allogrooming in cattle is currently unclear. Higher social dominance status has previously been found to be both positively (Šárová et al., 2016; Wood, 1977) and negatively (de Freslon et al., 2020) related to received allogrooming in cattle, as well as to have no relationship (Sato et al., 1993; Val-Laillet et al., 2009). Age-related characteristics (such as body mass, dominance, and the likelihood of having familiar group mates) may be linked to each other and create confounding effects on the rate of social behaviours. For example, cows which are more familiar with each other display higher allogrooming performance rates, with longer cohabitation increasing allogrooming frequency (Gutmann et al., 2015; Gygax et al., 2010), and older cows have been found to be more active groomers (de Freslon et al., 2020). The inconsistent relationship between dominance and allogrooming may also be due to variation in breed and group composition, since previous research has almost always been performed on farmed cattle with few exceptions (Reinhardt et al., 1986). There are limitations associated with farm environments as these are often single-sex and age-clustered groups. With frequent membership changes, farmed cattle are at a disadvantage when establishing long-term social bonds as they cannot make their own association choices (Marino & Allen, 2017). Therefore, it is advantageous to carry out behavioural research on free-roaming and feral cattle populations, as they may be more suitable for understanding the evolution of social behaviours.

The feral cattle of Hong Kong provide a unique opportunity to examine the social behaviour of free-ranging cattle with minimal human management (Pinkham et al., 2022; So & Dudgeon, 2020). Once used as draught animals, cattle were released in country parks after the decline of agricultural activities in the 1970’s, where they have continued to live and reproduce as free-ranging feral groups (AFCD 2013; Pinkham et al., 2022). Hong Kong feral cattle have high heterogeneity in colour and body type, being a unique mix of taurine and indicine genes, and are genetically distinct from other cattle populations (including *B. taurus* and *B. indicus*; Barbato et al., 2020; Perroux et al., in prep.). Groups of Hong Kong feral cattle have large variation in group size and sex ratios, and individuals live together for many years, having the ability to choose their own associations.

In this study, we investigated the relationship between sex, dominance status, and allogrooming in feral, free-ranging cattle in Hong Kong. We determined the dominance hierarchy by recording agonistic behaviours, and we recorded allogrooming interactions between identifiable individuals. Our main aims were to (1) assess the hierarchy of a feral cattle herd, (2) describe the occurrence and distribution of allogrooming within the herd, and (3) examine the relationship between allogrooming rate, sex, and dominance status. Based on these aims and the results of previous ungulate research, we predicted the following; (1) males would achieve higher dominance status than females (2), allogrooming would be directed towards body regions that individuals cannot reach via self-grooming (3) allogrooming would be unequally distributed in the group, with bouts not being entirely reciprocal, and (4) more dominant individuals would receive more allogrooming from others (i.e., allogrooming directed up the hierarchy).

## METHODS

### Study site and population

We carried out observations from 11 February 2022 to 27 May 2022 on one cattle herd in Sai Kung East Country Park, Hong Kong SAR, China. These months typically experience dry and mild weather with occasional high humidity (Hong Kong Observatory, 2022). Sai Kung East Country Park is a country park of 4,494 hectares as defined by the Agriculture and Fisheries Conservation Department (AFCD) of Hong Kong, consisting of country trails, reservoirs, woodland, coastland and villages. The study site where the cattle herd frequented was approximately 62 hectares (22°22’23.0”N, 114°20’07.5”E), covered by grassy vegetation with scattered trees and shrub. This location is accessible to the general public through roads and hiking trails passing through the site, and cattle at this location are habituated to human presence and traffic.

We collected data from a mixed-sex cattle herd, consisting of 35 males and 24 females (total = 59), with the number of individuals in the group varying from 47 to 56 cattle per observation day. Cattle have been present at this site since 2010. In 2013, 21 individuals were relocated to the study site from Lantau South Country Park (LCP, 2017), with at least 7 of these individuals being present during our data collection. Only adults were present in this herd, with individuals ranging approximately from 4 years to 15 years in age (Sai Kung Bovid Watch (SKBW), pers. comm.). No calves or juveniles were present at this location due to the cattle population contraception strategy enacted by the AFCD animal management team (AFCD, 2013).

The cattle fed on grass and other natural vegetation in the country park, and were provided with around 500-600kg of supplementary hay per week from a local citizen group. This amount of supplementary hay equated to approximately 22.5 –32.0 % of the herd’s estimated weekly nutritional requirement (Salah et al., 2014). Fresh water was available ad libitum from a reservoir next to the observation site.

We recorded the identities and sexes of the cattle present for each observation session. The animals were individually identified by ear tag number (as administered by the AFCD) or their distinctive physical appearance (e.g., coat colour, ear markings, horn presence, horn shape) if a tag was not present. To ensure accuracy of animal and behaviour identification, we spent two preliminary observation sessions (total 370 min) to ensure that observers could individually identify the cattle. We created an identification guide with all individual cattle images, identification, sex, and description of physical features.

### Behavioural observations

We carried out behavioural observations for 14 days from 0900 to 1500 hours, for an average of 228 ± 15.62 (SE) min observation per day. Observation sessions ranged from 60 min to 300 min, with a total observation time of 3200 min. We used continuous all-occurrence animal sampling methods (Altmann 1974) to identify cattle involved in dyadic interactions, and to record specified agonistic and affiliative interactions between individuals. In each session, three to four trained observers standing at least 20m away from the cattle conducted continuous observation. Each observer stood in a separate location to record interactions between individuals using binoculars and a stationary telescope. We recorded the date, time, and the identity of each social partner to ensure the same interaction was not duplicated.

### Allogrooming behaviour

We defined allogrooming behaviour as repeated licking movements by one animal on the body part of another (de Freslon et al., 2020; Laister et al., 2011; Val-Laillet et al., 2009). We continuously recorded the occurrence of allogrooming for the entire session. We recorded the identities of the performer (animal initiating the grooming and licking the other animal) and the receiver (animal receiving the grooming). We only included allogrooming bout duration in the analysis when the start and end times had been seen and the observation was separate from the previous bout by at least ten seconds (Laister et al., 2011). We recorded distinct grooming bouts if the roles of the performer and receiver were reversed, if the performer stopped grooming for ten seconds or longer, or if the performer switched to a different receiver (Laister et al., 2011). We classified the groomed body regions into nine categories: head, ears, neck, shoulders, flank, belly, back, legs and rear (Supplementary Material, Figure S1). We only analysed allogrooming body region data for complete allogrooming bouts with specified start and stop times.

### Agonistic behaviours

We recorded agonistic interactions within dyads which resulted in the receiver’s withdrawal from their original position by taking a minimum of 2 steps from the actor (Foris et al., 2021). Individuals were identified and assigned as winners (i.e. cattle who cause withdrawal) or losers (i.e. cattle which withdraw) depending on the outcome of the dominance interaction (Foris et al., 2021; Hubbard et al., 2021).

### Exclusion of individuals

We observed a total of 59 cattle over the observation period. Following visual inspection of the observation time histogram, we excluded animals which were present in the herd for less than 600 min of observation time. We removed the corresponding dyadic interactions from the interaction matrix in order to prevent biases towards those that were infrequently seen but active when they were present (Bray & Gilby, 2020). After exclusion of four individuals (2 males, 2 females), we used 55 cattle in the analyses (22 males, 33 females). We excluded interactions with unidentified individuals, missing data or those without a clear outcome from the analysis.

### Directional consistency and allogrooming exchange rates

We calculated the Directional Consistency Index (DCI) using package ‘EloRating’ to assess directionality in allogrooming bouts and in agonistic dominance interactions (Neumann & Kulik, 2020; Silk et al., 2013). In order to account for variable cattle presence in the group, we calculated relative rates of performing and receiving allogrooming per individual (Silk et al., 2013). For each individual, we created a relative time factor by dividing the number of minutes an animal was observed in the group by the total observation time at the group location. Then, the total number of allogrooming bouts performed or received by this individual (number of observations of the individual during the entire length of observation) were multiplied by this factor, hence creating individual relative rates of performing and receiving allogrooming.

### Calculation of the dominance hierarchy

The dominance status of each individual was determined through calculating a dominance hierarchy based on the direction of agonistic interactions. From dyadic interaction data, we created a win-lose agonistic interaction matrix for all agonistic interactions using R package ‘EloRating’ (Neumann & Kulik, 2020). Following guidelines outlined by Sánchez-Tójar et al. (2018), we calculated that the interaction-to-individual ratio and data sparseness were adequate to provide a reliable estimation of the hierarchy. To ensure a hierarchy was present and results have internal validity, the hierarchy was assessed for linearity, transitivity, stability, steepness and repeatability, using packages ‘EloRating’ (Neumann & Kulik, 2020) and ‘aniDom’ (Sánchez-Tójar & Farine, 2021). Additional comprehensive hierarchy assessment methodology is available within Supplementary Material, Information S1.

We used a training-testing procedure (R package ‘rankReliability’) to select the most reliable hierarchy ranking methodology with a data-splitting approach using 80% training data and 20% testing data (Vilette et al., 2020). We found that the randomised Elo-rating method developed by Sánchez-Tójar et al. (2018) had the highest reliability (0.89) to correctly predict the outcome of an interaction between two individuals (Supplementary Material, Information S1). This amended method of Elo-rating reduces temporal biases by randomising the interaction order by *n* times. Randomisations are then used to produce mean individual dominance scores. We applied the randomised Elo-rating method thereafter with 1000 randomisations to calculate mean hierarchy scores of each individual in the group, using R package ‘aniDom’ (Sánchez-Tójar & Farine, 2021). When not converted to ordinal ranks, higher randomised Elo-rating scores characterise highly dominant animals, with lower scores characterising subordinate animals (Sánchez-Tójar et al., 2018).

### General analyses and code availability

Statistical analyses were conducted with R (v4.2.0, R Core Team). We used a Chi-square goodness of fit test to determine whether allogrooming was randomly distributed on different body regions, using function ‘*chisq_test’* of package ‘stats’. We used Kendall correlation to test if individual allogrooming performance and recipient rates were correlated (function *‘cor.test’*, package ‘stats’). We also used an Analysis of Variance (ANOVA) model to test if males and females differed in dominance scores, using function ‘*aov*’ from package ‘stats’. Wherever ANOVA was used, homogeneity of the sample variances was tested using Bartlett tests (function ‘*bartlett.test’*, package ‘stats’), and residual normality was confirmed using Shapiro-Wilk tests (function ‘*shapiro.test’*, package ‘stats’). Wilcoxon tests were used to test whether the relative rate of allogrooming performance and receiving towards males or females differed between the sexes (function ‘*wilcox.test*’, package ‘stats’). Linear models were used to test if dominance status was related to receiving or performing allogrooming, and models were fitted using ‘*lm’* function of package ‘stats’ after we verified model assumptions using package ‘DHARMa’ (Hartig & Lohse, 2022). Where necessary to fulfil model assumptions, data were subject to log or square root transformation before analysis (Bateson & Martin, 2021). When transformations were not sufficient, rates were transformed into binary data of “Behaviour Present” (1) or “Behaviour Absent” (0), and ANOVA models were then used to test the effect of dominance status on the presence of the behaviour (function ‘*aov*’, package ‘stats’). R-code and data are available online at Hodgson et al., (2023).

### Ethical note

This research was reviewed and approved by the Animal Research Ethics Sub-Committee of City University of Hong Kong (Internal Reference: A-0826). We observed feral cattle from a 20m distance without any human intervention, and minimised disturbance to the animals in accordance with the Association for the Study of Animal Behaviour ethical guidelines (ASAB Ethical Committee & ABS Animal Care Committee, 2022).

## RESULTS

### Summary of dominance and allogrooming interactions

We recorded a total of 1155 agonistic dominance interactions. The mean number of dominance interactions per individual (including acting and receiving agonistic behaviours) was 42.00 ± 1.94 (SE), with a range of interactions per individual between 14 and 73. Dominance interactions were highly unidirectional and asymmetrical (Directional Consistency Index, DCI = 0.95), with lower-ranking individuals rarely displacing higher-ranking individuals.

We recorded a total of 422 allogrooming events. The mean number of allogrooming interactions (including both received and performed bouts) per individual was 15.35 ± 1.64 (SE), with allogrooming performance interactions ranging from 0 to 54 and receiving interactions ranging from 1 to 27. Relative rates of allogrooming performance ranged from 0 to 54, with cattle having a mean performance rate of 7.37 ± 1.27 (SE) over their observation time, and allogrooming receiving relative rates ranged from 0.70 to 28, with a mean receiving rate of 7.41 ± 0.74 (SE) over their observation time. Cattle allogroomed between 1 and 26 other individuals. The mean duration of complete allogrooming events was 72.80 ± 6.38s (SE) with complete bouts ranging from 1 to 550s. Overall, 46 individuals (19 males, 27 females) both performed and received allogrooming, while the remaining nine (3 males, 6 females) individuals received allogrooming but did not perform. Allogrooming exchanges were unidirectional and asymmetrical (Directional Consistency Index, DCI = 0.81), with bouts not being evenly exchanged between dyads, and one partner performing more allogrooming than the other.

### Body region location of allogrooming

Allogrooming body region data were recorded for 183 complete allogrooming bouts, with multiple regions being groomed during 92 events. Allogrooming across different body regions was not randomly distributed (chi-square test: χ^2^_8_ = 237.08, *P* < 0.001, Figure 1). The neck and head were the most commonly groomed regions, whereas fewer allogrooming events were performed on the legs and belly (Figure 1).

**Figure 1:**
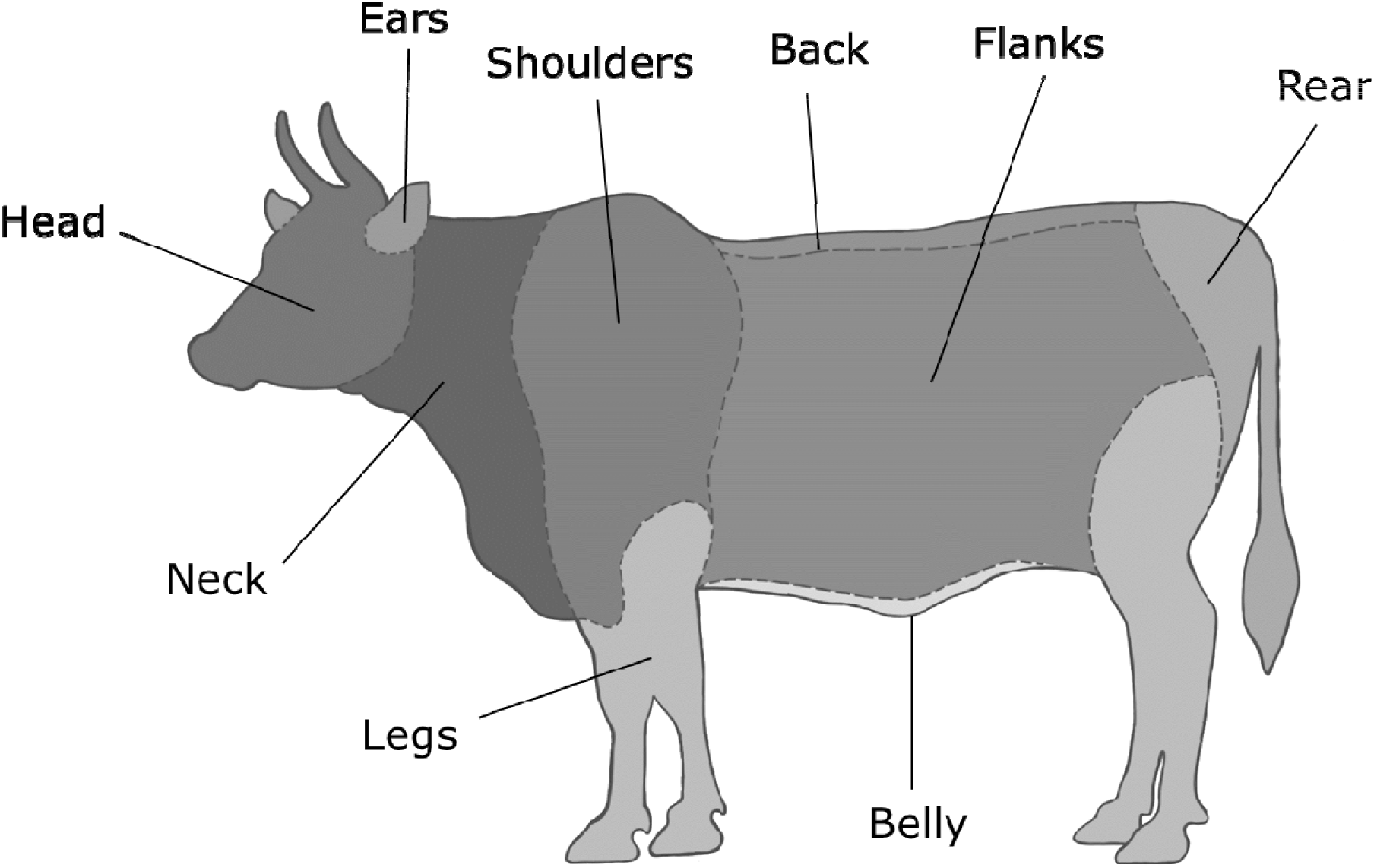
Allogrooming counts performed on different feral cattle body sites. Darker colour represents higher count frequency and lighter colour represents lower count frequency. From most to least allogroomed: neck (85), head (84), shoulders (51), flanks (39), back (36), ears (13), rear (12), legs (2) and belly (1).

### Effect of sex on dominance status

Sex affected the dominance score of cattle (ANOVA: *F*_1_ = 32.92, *P* < 0.001), with males having higher average dominance scores than females (Figure 2). However, not all female cattle were subordinate to male cattle, with one female in the top 20% of the ranking, and one male in the bottom 20% of the dominance hierarchy (Figure 2; Supplementary Material, Figure S2).

**Figure 2:**
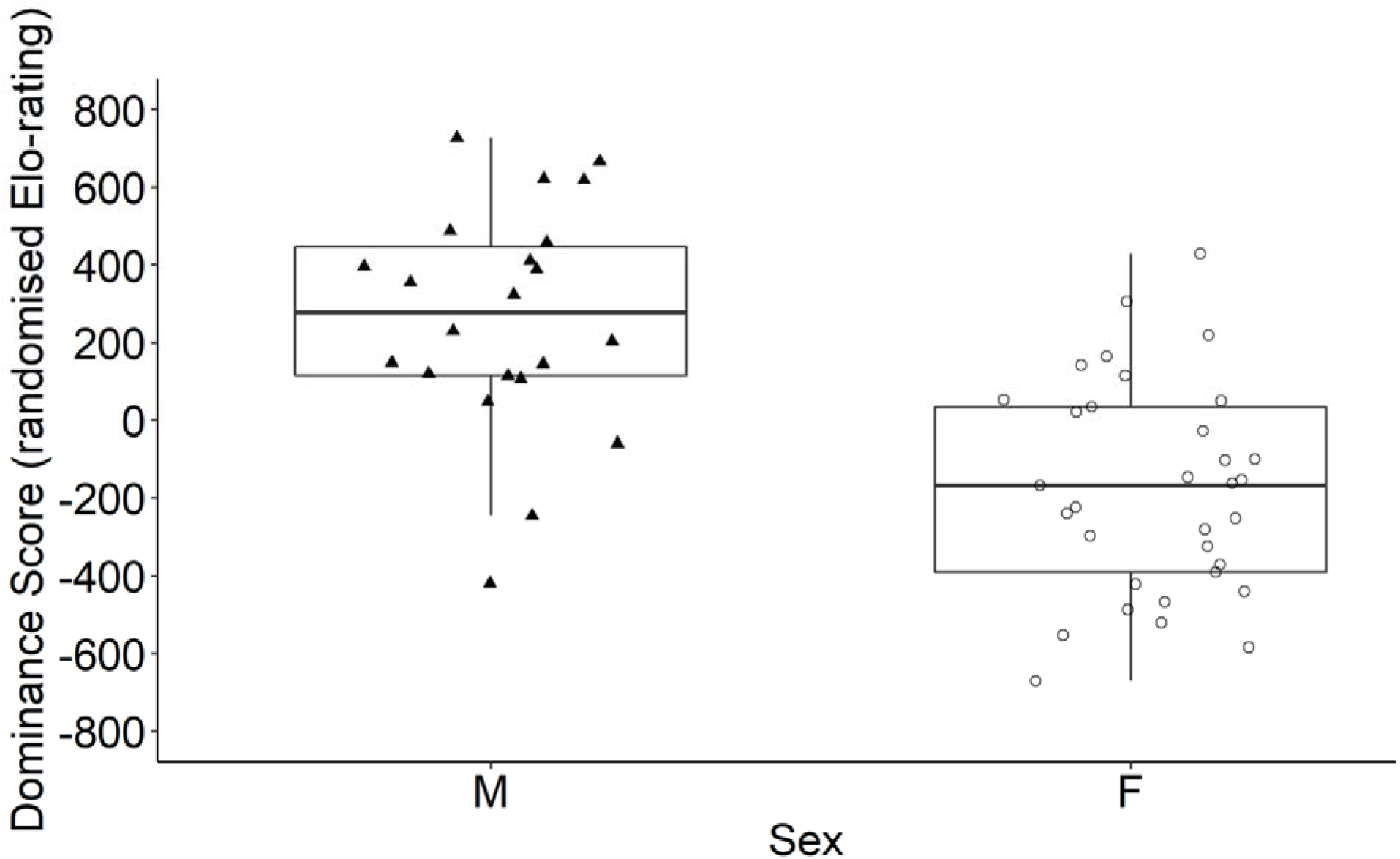
Average dominance scores of male cattle (**▴**; *n* = 22) and female cattle (**O**; *n* = 33). Dominance scores are calculated using agonistic interactions via randomised Elo-rating methodology. Dominance scores are expressed relative to the whole herd, with lower scores indicating subordinate animals in the hierarchy, and higher scores indicating dominant animals. Boxes indicate the first and third quartiles, with centres indicating the median values.

### Allogrooming performance and receiving rates

The relative individual rates of allogrooming performance and receiving were not related (Kendall correlation: τ = 0.23, *z* = 2.41, *P* < 0.05; Supplementary Material, Figure S3). This correlation meant that the likelihood of each individual performing allogrooming could not be used to predict their likelihood of being groomed by others, and similarly, receiving allogrooming could not predict an individual’s likelihood of performing it. This remained true for male performance and receiving (Kendall correlation: τ = 0.19, *z* = 1.17, *P* = 0.24) and for females (Kendall correlation: τ = 0.26, *z* = 2.08, *P* < 0.05) when analysed separately.

### Relationship between dominance and allogrooming

Dominance status affected the relative rate of receiving allogrooming, with higher-ranked individuals receiving more allogrooming (LM: R^2^ adj = 0.17, F_(2, 52)_ = 6.44, *P* < 0.05). However, this was only correct for higher-ranked females (LM: R^2^ adj = 0.30, F_(1, 31)_ = 14.98, *P <* 0.001; Figure 3), as higher-ranking males did not receive more (LM: R^2^ adj = 0.01, F_(1, 20)_ = 0.25, *P* = 0.28). There was no effect of dominance status on individual performance rates of allogrooming of neither males (LM: R^2^ adj = 0.07, F_(1, 20)_ = 2.60, *P* = 0.12) nor females (LM: R^2^ adj = -0.03, F_(1, 31)_ = 0.05, *P* = 0.82).

**Figure 3:**
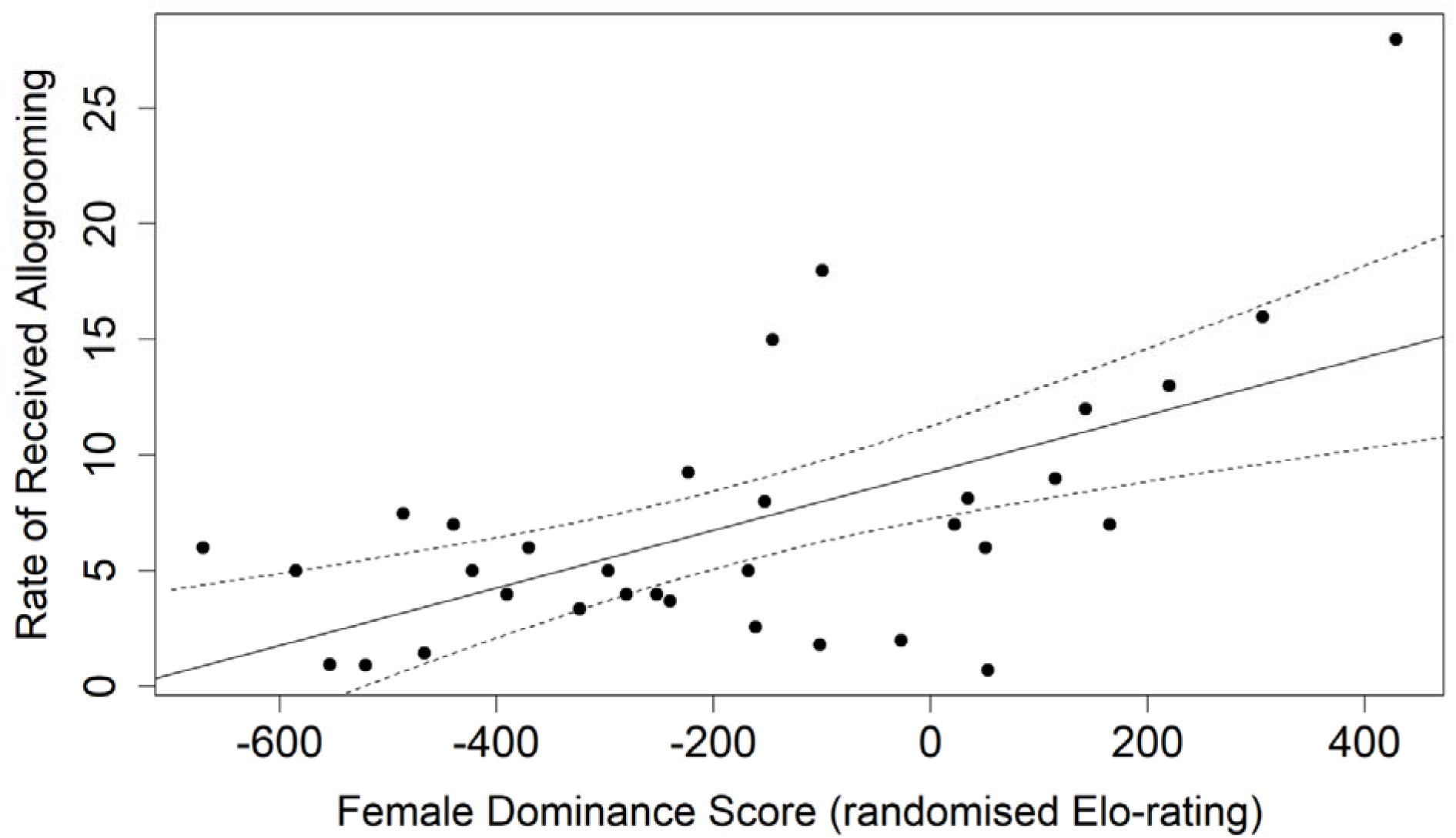
The relative rate of allogrooming received for female cattle (*n* = 33). Trend line and confidence intervals (95%) are from the linear regression model (LM: R^2^ adj = 0.30, F_(1, 31)_ = 14.98, *P <* 0.001). Dominance scores are calculated using agonistic interactions via randomised Elo-rating methodology. Dominance scores are expressed relative to the whole herd, with lower scores indicating more subordinate animals, and higher scores indicating more dominant animals.

### Effect of sex on individual allogrooming rates

Sex did not affect the rate at which an individual performed allogrooming (Kruskal-Wallis: χ^2^_1_ = 0.37, *p* = 0.53), nor the rate at which it received allogrooming (Kruskal-Wallis: χ^2^_1_ = 0.79, *P* = 0.38). Sex did not affect the number of allogrooming partners an animal had (Kruskal-Wallis: χ^2^_1_ = 0.59, *p* = 0.44). After separating allogrooming performance rates towards all individuals into separate allogrooming performance rates towards males and towards females, males performed more allogrooming bouts towards females than other males (Wilcoxon signed-rank test: *V* = 25.50, *n* = 22, *P* < 0.05; Figure 4). Females performed allogrooming equally towards both males and other females (Wilcoxon signed-rank test: *V* = 139.00, *n* = 33, *P* = 0.76). Males received equal amounts of grooming from other males and from females (Wilcoxon signed-rank test: *V* = 92.50, *n* = 22, *P* = 0.43), and females received equal grooming from males and other females (Wilcoxon signed-rank test: *V* = 305.00, *n* = 33, *P* = 0.27).

**Figure 4:**
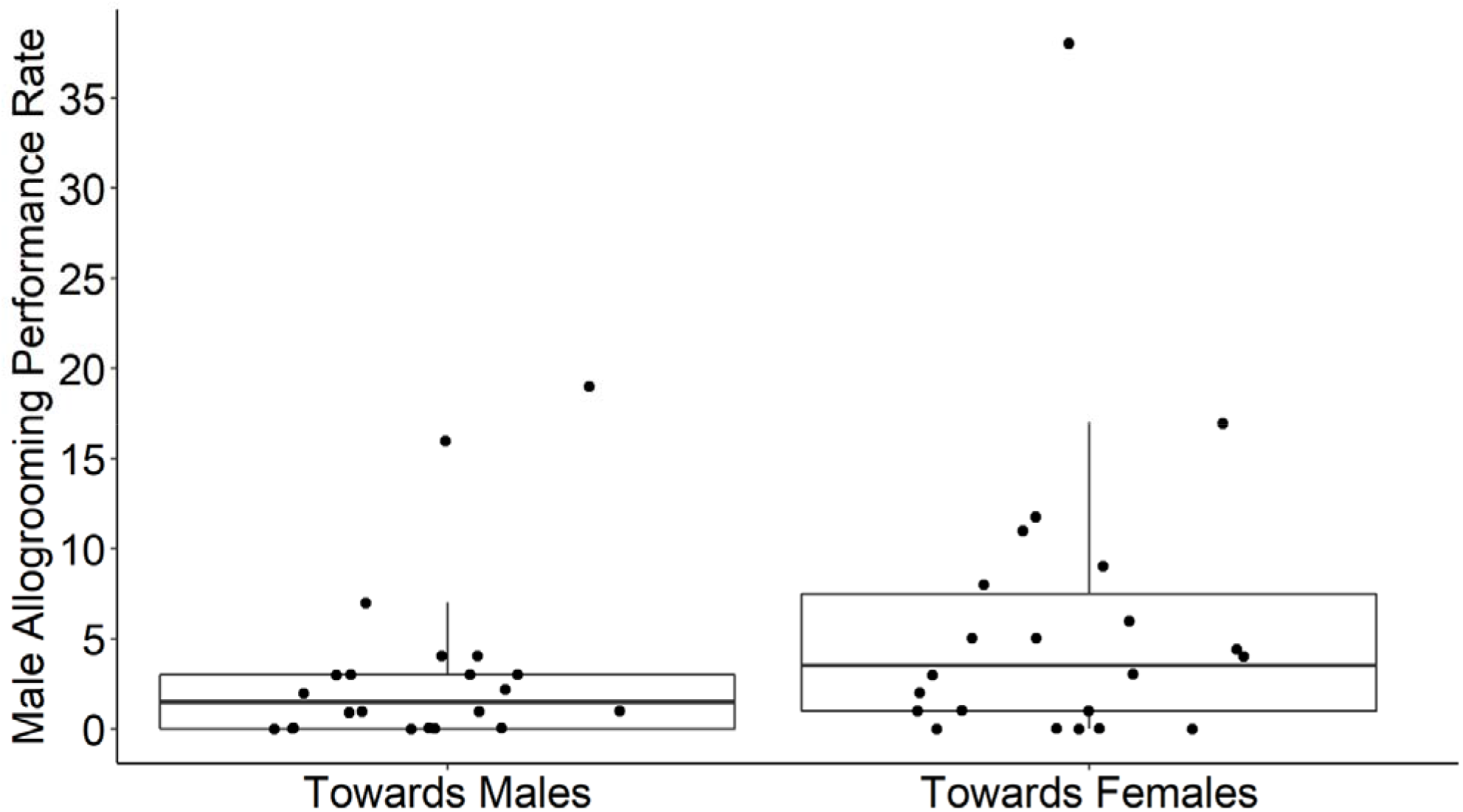
The relative rate of individual male cattle allogrooming performance (*n* = 22) towards males and towards females. Boxes indicate the first and third quartiles, with centres indicating the median values.

### Effect of dominance on sex-specific allogrooming

Female dominance status did not affect allogrooming performance rate towards other females (LM: R^2^ adj = 0.01, F_(1, 31)_ = 1.26, *P* = 0.27; Supplementary Material, Figure S4). Dominance score also did not affect the likelihood that a female would perform allogrooming towards a male (ANOVA: *F*_1_ = 0.09, *P* = 0.77). Females with higher dominance scores received more allogrooming from other females (LM: R^2^ adj = 0.10, F_(1, 31)_ = 4.38, *P* < 0.05), and from males (LM: R^2^ adj = 0.33, F_(1, 31)_ = 16.92, *P <* 0.001; Figure 5, Supplementary Material, Figures S5 and S6).

**Figure 5:**
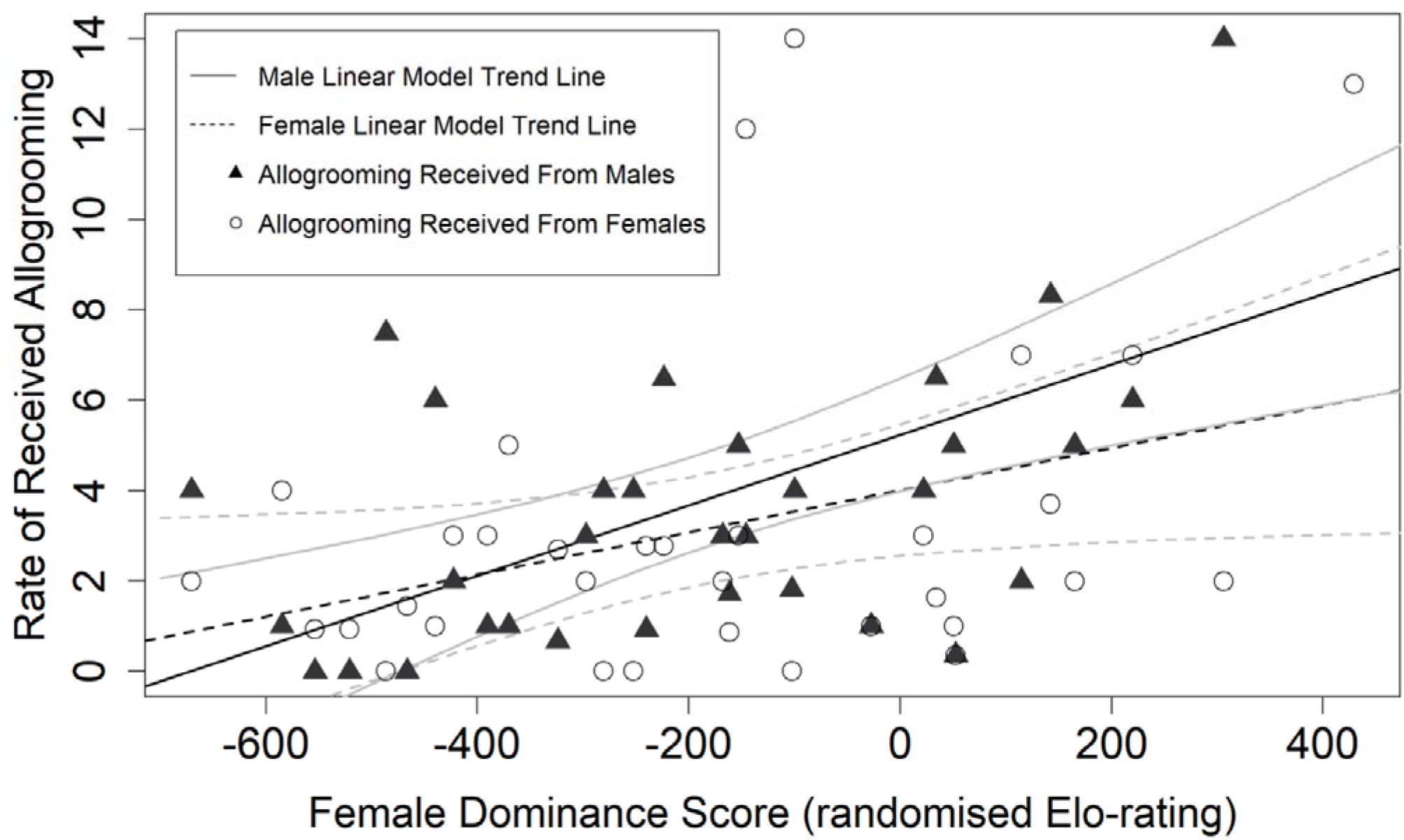
Distribution of allogrooming received from males (**▴**) and females (**O**) for female feral cattle. Linear regression trend lines and confidence intervals (95%) are from separate models; allogrooming received from females (LM: R^2^ adj = 0.10, F_(1, 31)_ = 4.38, *P* < 0.05), and received from males (LM: R^2^ adj = 0.33, F_(1, 31)_ = 16.92, *P <* 0.001).

There was a trend towards higher-ranking males performing more allogrooming towards females (LM: R^2^ adj = 0.11, F_(1, 20)_ = 3.60, *P* = 0.07), but dominance score did not affect the rate of male allogrooming towards other males (LM: R^2^ adj = -0.01, F_(1, 20)_ = 0.85, *P* = 0.37). Higher ranking males did not receive more allogrooming from females (LM: R^2^ adj = -0.01, F_(1, 20)_ = 0.72, *P* = 0.41), nor from other males (LM: R^2^ adj = -0.04, F_(1, 20)_ = 0.24, *P* = 0.63).

## DISCUSSION

Affiliative behaviours such as allogrooming are important for the survival, reproduction, and health of group-living animals, allowing us to understand how individuals interact with and respond to each other (Barocas et al., 2011; Bond et al., 2021; Riehl & Strong, 2018; Silk, 2007; Silk et al., 2003). We investigated the occurrence of allogrooming in relation to sex and dominance in feral cattle. We found that higher-ranking females received more allogrooming than lower-ranking ones, but by contrast, we did not find any effect of dominance on the likelihood of performing allogrooming. Our results show that males performed more allogrooming towards females than towards other males. Allogrooming was unevenly distributed on body regions, and frequently directed towards body regions that were inaccessible for the receiver to self-lick, such as the head or neck. We also found that individual allogrooming performance and receiving rates were not related. Our results show that males achieved generally higher dominance status than females, but not all females were subordinate to males. These sex-specific effects of allogrooming and dominance are critical to our understanding of sociality and the evolution of ungulate social behaviour, especially when under free-ranging conditions (Bowyer et al., 2020; Hubbard et al., 2021; Vander Wal et al., 2012).

### Dominance, sex & allogrooming

We found that higher-ranking females received more allogrooming from both males and females. Dominant individuals may receive more allogrooming and affiliative behaviours in exchange for providing services such as food or mates, as found in non-human primates such as chimpanzees (Feldblum et al., 2021) and macaques (Bhattacharjee et al., 2023). Allogrooming may also be strategically directed at dominant animals to increase appeasement, social tolerance, and reconciliation after conflict, as seen in meerkats (*Suricata suricatta*; Kutsukake & Clutton-Brock 2010), and vervet monkeys (*Chlorocebus pygerythrus*; Gareta García et al., 2021). Lower-ranking individuals can gain these rank-related benefits from their higher-ranking partners by directing allogrooming up the hierarchy (Gumert, 2007; Seyfarth, 1977). However, we did not find this typical pattern in feral cattle, as although higher-ranking females received more, both subordinate and dominant females performed equally. Previously, higher-ranking cattle have been found to exchange preferential allogrooming, suggesting that certain sub-groups may exist within the group (Šárová et al., 2016). For example, higher-ranking animals may only exchange allogrooming bouts within close ranks, whereas lower-ranking cattle may direct their grooming indiscriminately or only to higher ranking individuals. Higher-ranking females may also receive more allogrooming if they possess certain individual characteristics such as being older or more familiar, as both age and familiarity influence dominance in cattle and can also be positively related to received allogrooming (de Freslon et al., 2020; Hubbard et al., 2021; Šárová et al., 2013). We are unable to exclude the possibility that allogrooming is directed to older individuals, as we do not have exact ages or association duration for our study animals; however, age may explain differences in previous cattle research results (de Freslon et al., 2020; Sato et al., 1993; Val-Laillet et al., 2009; Wood, 1977). As Hong Kong cattle are a genetically distinct population (Barbato et al., 2020), differences in our results from previous research could be due to variation in evolutionary pressures and domestication processes (Cooke et al., 2020), with indicine and taurine cattle breeds differing in physiology (Sartori et al., 2016) and social behaviours (Solano et al., 2005).

We found that males performed more allogrooming towards females than other males. With a 2:3 male-to-female group sex ratio we cannot rule out the possibility that this bias is due to individuals being more likely to allogroom a female. However, this is unlikely to be the case, as female cattle were found to perform equally to both males and other females. Inter-sexual relationships are not frequently analysed in farmed ungulates due to husbandry constraints, with cattle predominantly kept in single-sex groups, but male and female cattle display a variety of sexually-specific behaviours (Solano et al., 2005). Our data from Hong Kong cattle provide a novel opportunity to examine these relationships, and feral mixed-sex herds could be used to better understand this link between sex and allogrooming. Although sex has not previously been found to affect allogrooming in cattle (Sato, 1984), sexually-dimorphic allogrooming occurs if the benefits of allogrooming are sex-specific, such as male vervet monkeys being rewarded with allogrooming by females after conflict participation (Arseneau-Robar et al., 2016). Males may preferentially allogroom females if it plays a courtship role to increase their reproductive success. For example, male singing mice (*Scotinomys teguina*) use allogrooming to secure reproductive access to females (Fernández-Vargas et al., 2011), and allogrooming can be exchanged for mating opportunities, as found in male white-handed gibbons (*Hylobates lar*; Barelli et al., 2011) and male long-tailed macaques (*Macaca fascicularis*; Gumert 2007). We suggest further investigation into male allogrooming performance rates to help identify sexually-specific social behaviours which contribute to group sociality.

### Distribution of allogrooming

Our results show that allogrooming was primarily directed towards inaccessible body regions such as the head and neck, whereas the least frequently allogroomed areas were those that the animal could self-lick. Although allogrooming plays a social function in many group-living species (Di Bitetti, 1997; Kutsukake & Clutton-Brock, 2010), it evolved to serve a key health benefit, helping to remove dirt particles and ectoparasites from skin and fur (Akinyi et al., 2013; Dunbar, 1991; Mooring et al., 1996). This hygienic purpose is commonly found among mammals, with cattle (Tresoldi et al., 2015; Val-Laillet et al., 2009), black capuchin monkeys (*Sapajus nigritus*; Pfoh et al., 2021) and horses (*Equus ferus caballus*; King & Gurnell 2019) all directing allogrooming towards inaccessible regions, consistent with our results and the hygienic purpose of allogrooming.

Allogrooming was not equally distributed, and we found no relationship between the individual rates of performance and receiving, suggesting that feral cattle do not use reciprocal allogrooming to maintain affiliative patterns. In other species, mutual affiliative behaviours between individuals help social bonds to form within stable groups (Alexander, 1974; Trivers, 1971). For example, simultaneous exchanges of grooming in horses (Cameron et al., 2009) and chimpanzees (Allanic et al., 2021), allow allogrooming to be exchanged without a time delay. However, similarly to previous research in farm cattle (Šárová et al., 2016; Sato, 1984), we find that all animals received allogrooming, but not all performed. In a previous farm study, some cattle dyads did not exchange allogrooming, with up to a fifth of partners never found to allogroom together (Val-Laillet et al., 2009). Benefits may be especially hard to identify if there is a time delay between allogrooming exchanges. Although this requires repeated interactions and bookkeeping of other group members (Stevens et al., 2005), cattle can easily discriminate between conspecifics (Nawroth et al., 2019), and have long-term memory of social partners (Gutmann et al., 2015). Trading allogrooming as a commodity in a biological market can affect the pay-offs of allogrooming and its distribution (Hammerstein & Noë, 2016; Schino & Aureli, 2009). This means that other services (such as tolerance while feeding or shade acquisition) could be being exchanged for grooming without a reciprocal pattern being evident, and the distribution of desirable commodities should be carefully examined to evaluate if reciprocal trading is occurring (Barrett et al., 1999).

### Sex and dominance

We found that sex affected dominance score, with males ranked higher in the hierarchy. However, we also found that males were not always dominant over all females. Cattle are sexually dimorphic ungulates, with males displaying larger body masses than females (Doyle et al., 2021; Perroux et al., in prep.; Polák & Frynta, 2010). In many species, body mass determines an individual’s competitive ability and the likelihood of winning conflict (Arnott & Elwood, 2009; Clutton-Brock, 2017). Body mass is positively related to dominance rank in other dimorphic ungulates, such as fallow deer (*Dama dama;* McElligott et al., 2001) and bighorn rams (*Ovis canadensis*; Pelletier & Festa-Bianchet 2006). In certain species, dominance is sex-independent and both females and males attain similar dominance ranks, such as rock hyrax (*Procavia capensis*; Koren et al., 2006) and bonobos (*Pan paniscus*; Surbeck & Hohmann 2013). Although uncommon, hierarchies can also be female-biased; for example, males are consistently subordinate to females in sifaka (*Propithecus verreauxi*; Kappeler & Schäffler 2008) and spotted hyenas (*Crocuta crocuta*; Goymann et al., 2001). Among primates, the likelihood of female-biased hierarchies also depends on group composition; female dominance is found to be greater in groups with higher aggression and those with a higher proportion of males (Hemelrijk et al., 2008). As our study was conducted on a group with an approximately 2:3 male-to-female sex ratio, this is a potentially interesting future approach to studies with feral animals and comparisons between different ungulate species.

Dominance is affected by traits other than body size, such as age, affiliative interactions, parentage, and personality (Bagnato et al., 2023; Goymann et al., 2001; Holekamp & Smale, 1991; Schino, 2007; Schülke et al., 2010). Age affects dominance in cattle, with older animals dominating younger ones (Šárová et al., 2013). These effects can also be inter-sexual, with older females overruling males that are younger until a certain point (Hall, 1986; Hubbard et al., 2021; Reinhardt et al., 1986). However, all cattle in our study were adults, with probable age ranges from 4 years to 15 years, and as our group hierarchy linearity and transitivity were high, we assume that all dominance relationships had been well-established.

### Conclusion

Overall, our research on allogrooming in feral cattle offers a unique insight into the relationship between ungulate sex, dominance, and affiliative behaviour. We found that higher-ranking females received more allogrooming from other individuals, but lower-ranking animals did not perform more. Males allogroomed females more than other males, suggesting a sex-specific aspect to cattle behaviour that is not usually evident when studying the species on farms. We suggest that grooming is not directed up the hierarchy to exchange rank-related benefits as found in primates, but may instead be used to strengthen social bonds and promote affiliation within the group. These results point to the importance of agonistic and affiliative behaviours in social relationships, especially in sexually dimorphic species such as cattle. Our findings improve our understanding of allogrooming in mixed-sex stable social groups, and reinforce the use of allogrooming as a sociality indicator in ungulates.

## ACKNOWLEDGEMENTS

We are grateful to Sai Kung Bovid Watch, Hong Kong Bovid Conservation Association, and the AFCD cattle team for providing valuable information on our study animals. We thank Claire Giraudet for help with data collection, and Dr. Debottam Bhattacharjee for his valuable feedback on an earlier version of this manuscript.

## AUTHOR CONTRIBUTIONS

AGM, KJF, GMWH and TAP conceptualised the study. GMWH, TAP, WYC performed data collection in the field, data analysis, and drafted the original manuscript. GMWH, TAP, WYC, KJF and AGM critically reviewed and edited the manuscript. All authors have read and approved the final version of the manuscript and agree to be held accountable for the content of this paper.

