## Supplementary Information for "Sex and dominance status affect allogrooming in free-ranging feral cattle"

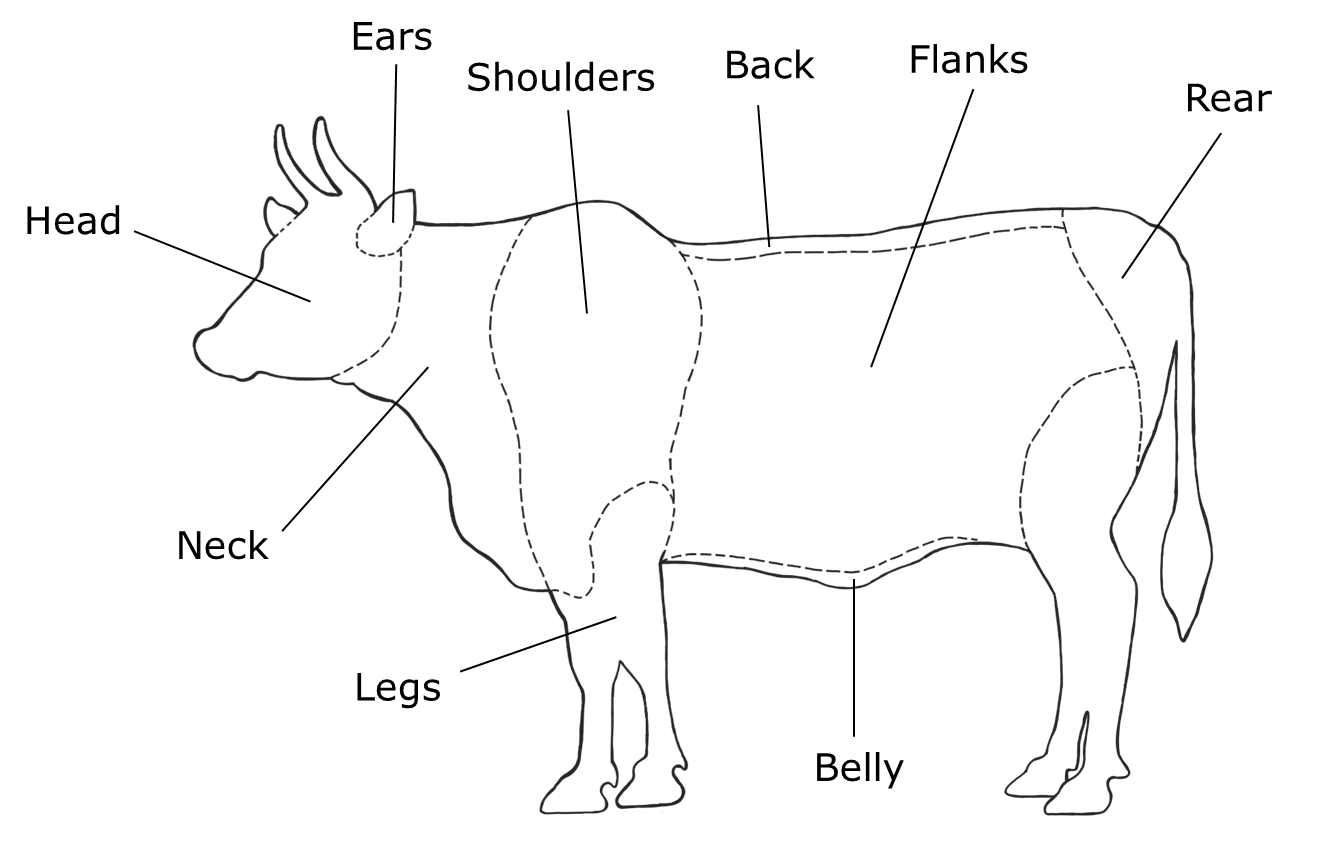

**Supplementary Material, Figure S1: Differentiation of cattle body regions; head (head region rostral to the jawline, excluding the ears), ear (ear pinna, only when it could be clearly seen that tongue was in contact with inner concave or outer convex surface), neck (including dewlap; from back of head and jawline to shoulder joint), shoulders (region dorsal to and including the shoulder joint of forelimbs), flank (lateral side of the body), belly (ventral part of the body), back (from shoulders to the start of the tail base; dorsal part of the body), legs (forelimbs and hindlimbs of the animal), rear (from tip of the tail to the tail base and hip joint, including genitalia).**

### SUPPLEMENTARY MATERIAL, INFORMATION S1

Although many methods for establishing a dominance order from observational data exist, there are two main families of ranking methodology; sequential, when the interaction sequence is accounted for, and non-sequential, when the order of interactions does not matter and the method instead relies on interaction matrices. Both have been found to have similar predictions, with neither family yielding more reliable results, and the best method for inferring a hierarchy may instead be specific to each dataset (Vilette et al., 2020). In order to select the method which provides the most accurate hierarchy for our dataset, we followed the data-splitting approach outlined in Vilette et al., 2020, which allows different hierarchies to be compared in prediction accuracy. We compared David’s Score (via R package ‘EloRating’; (David, 1987; Neumann & Kulik, 2020), Elo-Rating through two different methods (the first Elo-Rating v.1 through R package ‘aniDom’ and the second Elo-Rating v.2 via R package EloRating; Neumann and Kulik 2020; Sanchez-Tojar 2021), and Randomised Elo-Rating (via R package aniDom; Sanchez-Tojar 2021). Hierarchies were created based on 80% of the data, then tested against the remaining 20% of data to assess which hierarchy most reliably predicted the outcomes.

Reliability was high for all but estimated to be highest for the Randomised Elo-Rating methodology (0.887), followed by Elo-Rating v.2 (0.866), Elo-Rating v.1 (0.853), and David’s Score (0.848). As it provided the most accurate prediction of the outcome of an interaction between two individuals, the Randomised Elo-rating was used thereafter to calculate the hierarchy. This method is a non-sequential variation of the original Elo-Rating method. As scores generated from the original Elo-Rating are affected by the temporal sequence of the data, scores can be biased towards periods of high or low interaction. This method reduces potential biases causes by temporal variation by using the mean Elo-scores for each individual, calculated from 1000 randomisations of the sequence interaction data.

*Supplementary Hierarchy Assessment Methodology:*

We calculated stability through function ‘*stab_elo’*, with index values ranging from zero to one, where one indicates a stable hierarchy with no rank changes and zero indicates an unstable hierarchy with many rank changes. We estimated linearity via Landau’s linearity index (*h’*) using 1000 permutations and function ‘*h.index* (de Vries 1995). However, as Landau’s *h’* is negatively biased when datasets have an increased number of unknown relationships, we also calculated linearity using the triangle transitivity index (TTRI) as a measure of dominance orderliness through function ‘*transitivity*’ (Shizuka and McDonald 2012; Sánchez-Tójar et al. 2018). TTRI values range from 0 (when transitive relationships are random in a population) to 1 (where all triangle relationships are transitive). We calculated the steepness of the hierarchy based on adjusted, normalised David’s Scores using function ‘*steepness*’ (de Vries et al. 2006). The steepness of a hierarchy allows us to infer how adjacently ranked individuals differ in their success at winning dominance interactions, as large differences lead to a steeper hierarchy, and small differences indicate a shallow hierarchy (de Vries et al. 2006).

We calculated uncertainty of the hierarchy using two methods within R package ‘aniDom’; uncertainty estimated by repeatability of individual randomized scores via function ‘*estimate_uncertainty_by_repeatability’*, and uncertainty estimated by the correlation score (*r*_S_) between two inferred hierarchies after data-splitting via function ‘*estimate_uncertainty_by_splitting’* (Sanchez-Tojar 2021). Repeatability scores above 0.65 and correlation scores above 0.5 indicate that a hierarchy is certain and steep; repeatability and correlation scores above 0.9 indicate a highly certain and very steep hierarchy (Sánchez-Tójar et al. 2018).

#### *Hierarchy Structure*

The group hierarchy was significantly linear (h’ = 0. 2195, h expected = 0.0531, *p =* 0.001), highly transitive (Pt = 0.985, ttri = 0.938, *p* <0.001), and highly stable (stability = 0.986). The hierarchy was significantly steep (steepness = 0.125, expected 0.0391, *p* = 0.01) and repeatable, with a correlation score between inferred hierarchies of 0.865 (2.5% CI = 0.813, -97.5% CI = 0.914) and repeatability score of 0.929. Together, these values indicate that the hierarchy had low uncertainty and was more ordered than expected by chance.

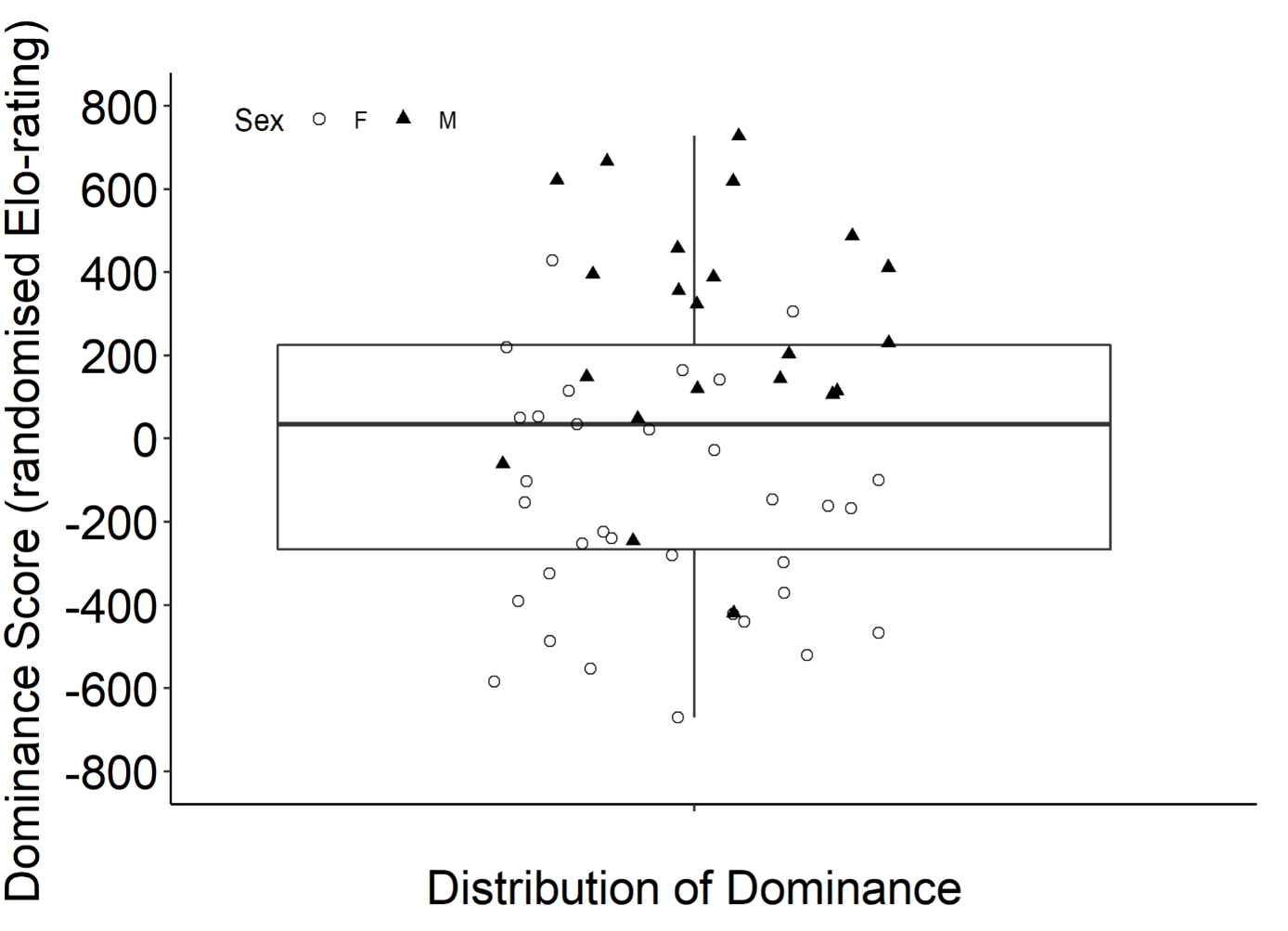

**Supplementary Material, Figure S2: Distribution of dominance scores (calculated using agonistic interactions via randomised Elo-rating methodology) in the herd hierarchy, including females (**O**)** **and males (▲). Boxes indicate the first and third quartiles, with centres indicating the median values.**

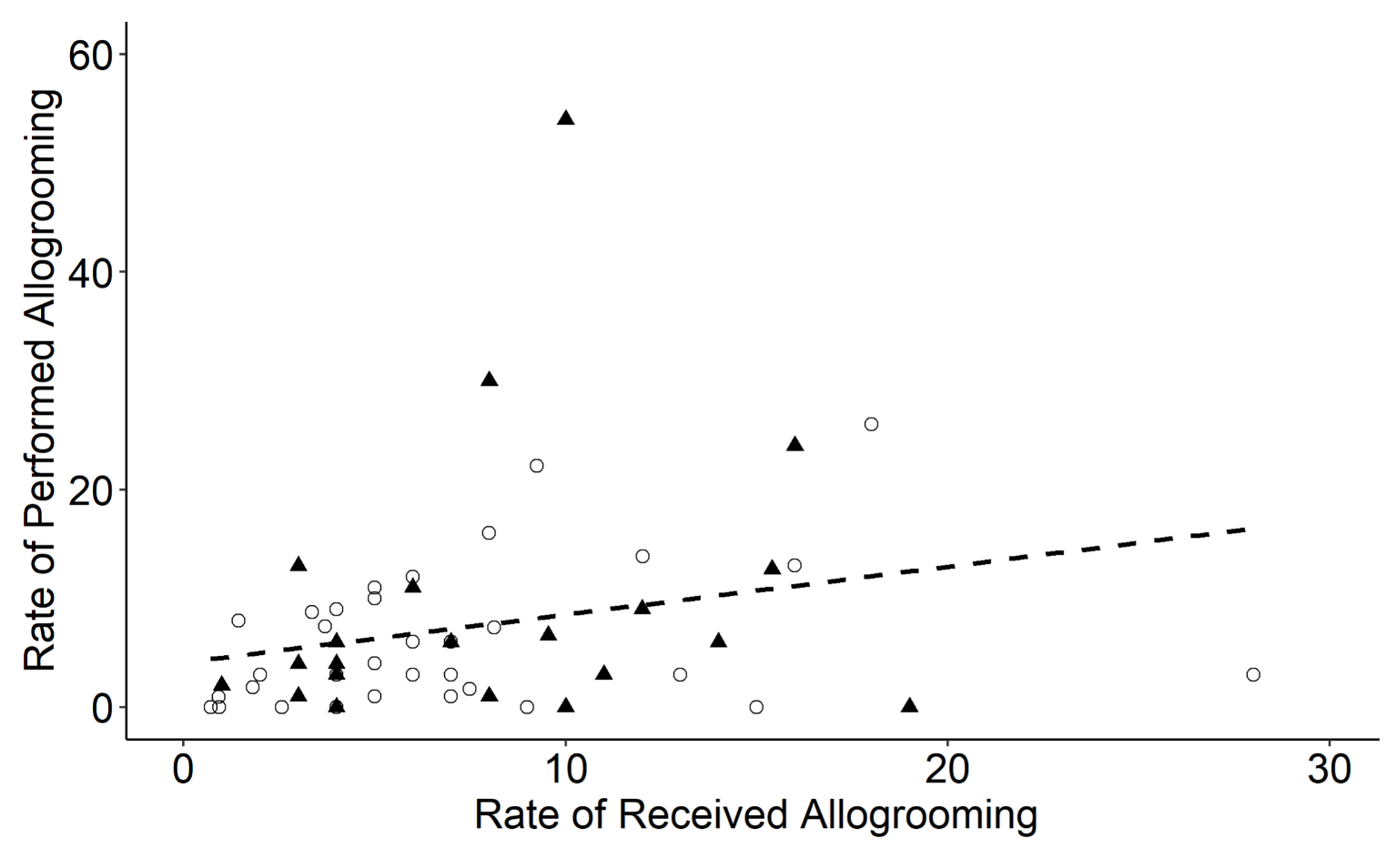

**Supplementary Material, Figure S3: Relative rates of allogrooming for individual cattle in feral cattle, including males (▲) and females (**O**), using trend line based on linear regression.**

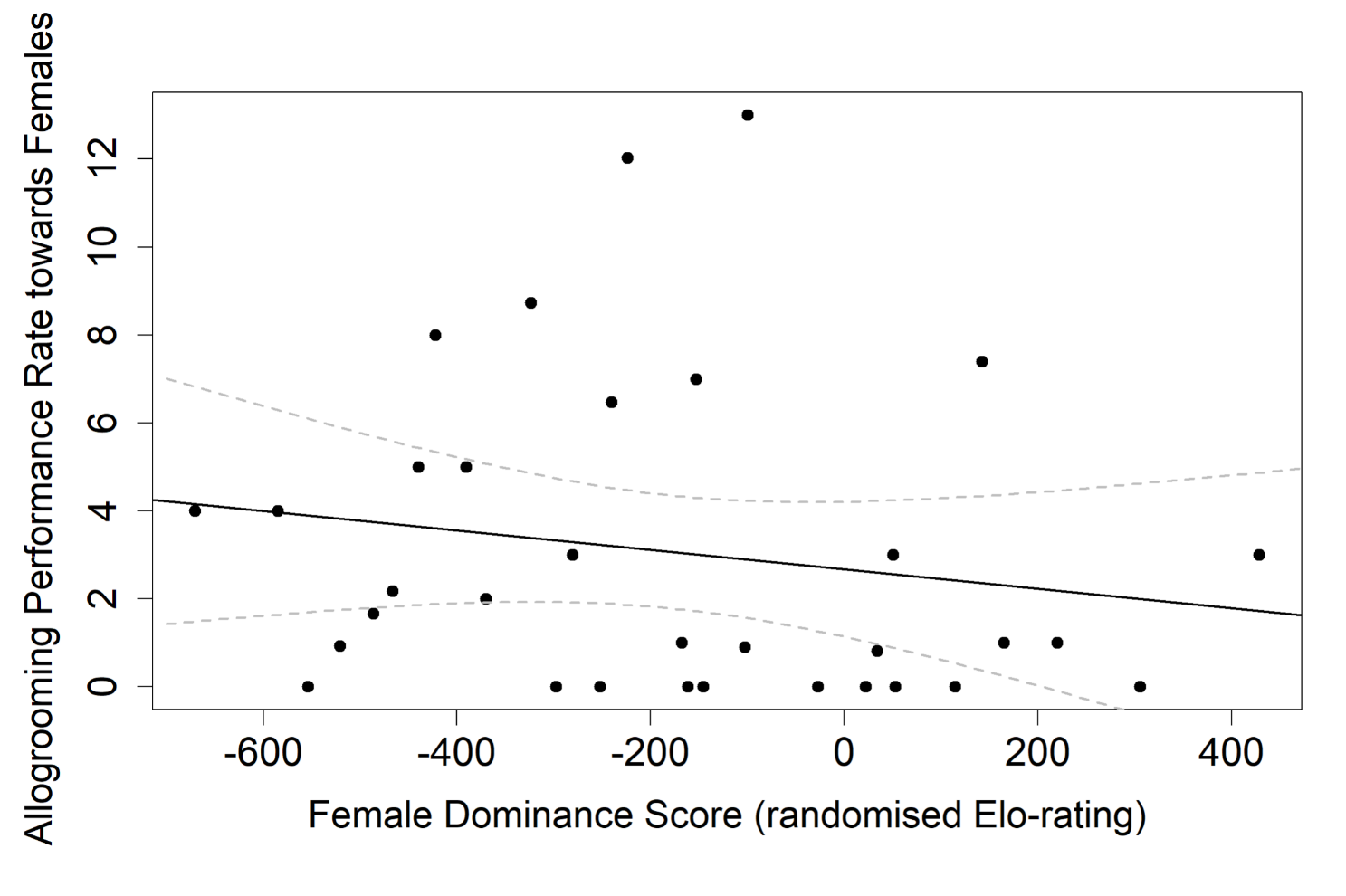

**Supplementary Material, Figure S4: Distribution of female allogrooming performance towards other females in the herd. Regression line and confidence intervals (95%) calculated from linear model (LM: R^2^ adj = 0.01, F_(1, 31)_ = 1.26, *P* = 0.27). Dominance scores are calculated using agonistic interactions via randomised Elo-rating methodology, where lower scores indicate subordinate animals in the hierarchy, and higher scores indicate dominant animals.**

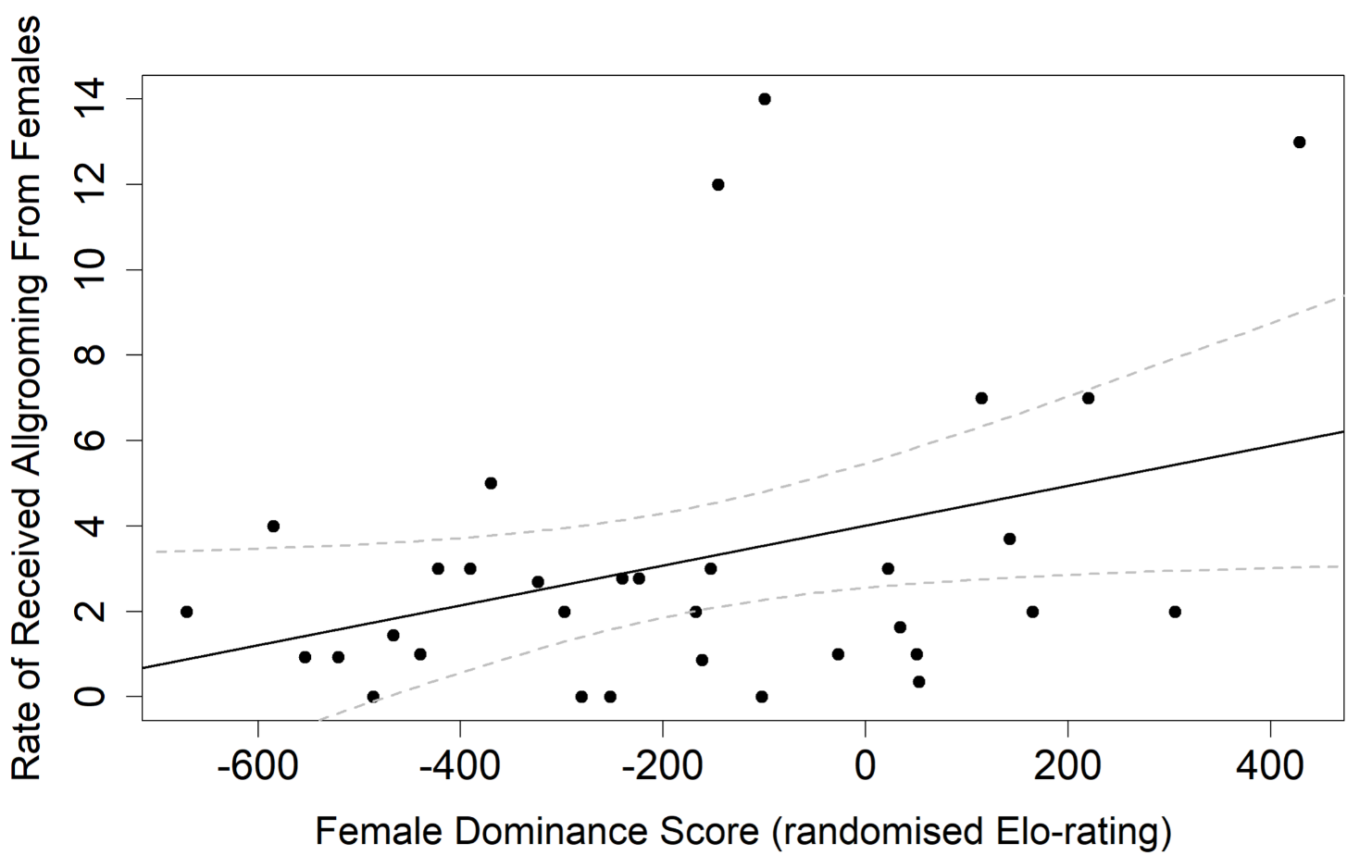

**Supplementary Material, Figure S5: Distribution of the relative rate of allogrooming received from other females. Regression line and confidence intervals (95%) calculated from linear model (LM: R^2^ adj = 0.10, F_(1, 31)_ = 4.38, *P* < 0.05). Dominance scores are calculated using agonistic interactions via randomised Elo-rating methodology, where lower scores indicate subordinate animals in the hierarchy, and higher scores indicate dominant animals.**

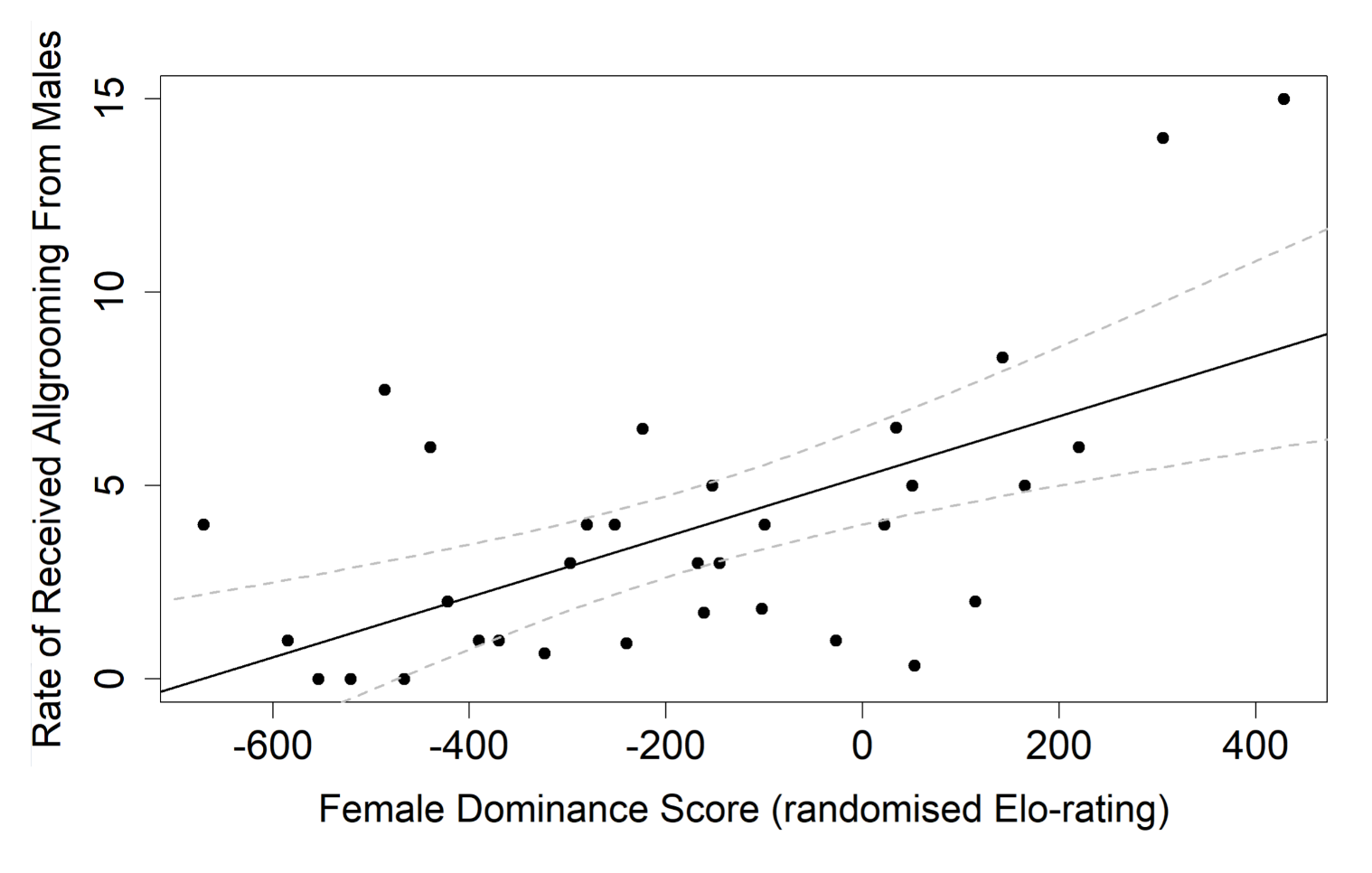

**Supplementary Material, Figure S6: Distribution of the relative rate of allogrooming received from males. Regression line and confidence intervals (95%) calculated from linear model (LM: R^2^ adj = 0.33, F_(1, 31)_ = 16.92, *P <* 0.001). Dominance scores are calculated using agonistic interactions via randomised Elo-rating methodology, where lower scores indicate subordinate animals in the hierarchy, and higher scores indicate dominant animals.**
